# Multicomponent depolymerization of actin filament pointed ends by cofilin and cyclase-associated protein depends upon filament age

**DOI:** 10.1101/2024.04.15.589566

**Authors:** Ekram M. Towsif, Blake Andrew Miller, Heidi Ulrichs, Shashank Shekhar

## Abstract

Intracellular actin networks assemble through the addition of ATP-actin subunits at the growing barbed ends of actin filaments. This is followed by “aging” of the filament via ATP hydrolysis and subsequent phosphate release. Aged ADP-actin subunits thus “treadmill” through the filament before being released back into the cytoplasmic monomer pool as a result of depolymerization at filament pointed ends. The necessity for aging before filament disassembly is reinforced by preferential binding of cofilin to aged ADP-actin subunits over newly-assembled ADP-P_i_ actin subunits in the filament. Consequently, investigations into how cofilin influences pointed-end depolymerization have, thus far, focused exclusively on aged ADP-actin filaments. Using microfluidics-assisted Total Internal Reflection Fluorescence (mf-TIRF) microscopy, we reveal that, similar to their effects on ADP filaments, cofilin and cyclase-associated protein (CAP) also promote pointed-end depolymerization of ADP-P_i_ filaments. Interestingly, the maximal rates of ADP-P_i_ filament depolymerization by CAP and cofilin together remain approximately 20–40 times lower than for ADP filaments. Further, we find that the promotion of ADP-P_i_ pointed-end depolymerization is conserved for all three mammalian cofilin isoforms. Taken together, the mechanisms presented here open the possibility of newly-assembled actin filaments being directly disassembled from their pointed-ends, thus bypassing the slow step of P_i_ release in the aging process.

## Introduction

Cellular actin dynamics are essential in a number of key processes such as cell migration, wound healing, cell division, and endocytosis (Carlier and Shekhar, 2017; Shekhar et al., 2016). Cells rapidly assemble and remodel their actin cytoskeleton to move, change shape, and reorganize their organelles. Intracellular actin networks consist of actin filaments, which assemble by polymerization of ATP-actin subunits at their barbed ends (Lappalainen et al., 2022). Upon addition of the subunit to the filament, the ATP bound to the actin subunit rapidly hydrolyzes. This results in the gamma phosphate being cleaved and the ATP-actin subunit getting converted into an ADP-P_i_-actin subunit (Blanchoin and Pollard, 2002). Rapid ATP hydrolysis is followed by a much slower step (0.0045 to 0.006 s^-1^), during which the cleaved phosphate is released, and the subunit converts to ADP-actin (Carlier, 1987; Carlier and Pantaloni, 1986). This ‘aging’ of actin subunits primes the filament for targeting by disassembly factors, such as cofilin, which have a much higher affinity for ADP-actin filaments than for ATP or ADP-P_i_ filaments (Blanchoin and Pollard, 1999). Consequently, cofilin-mediated disassembly of purified actin filaments is expected to be dictated by the age of actin filaments (Carlier et al., 1997; Maciver et al., 1991). Importantly, while actin filaments *in vitro* take tens of minutes to (age and) depolymerize, actin networks *in vivo* get turned over orders of magnitude faster. For example, actin filaments in the lamellipodia and endocytic patches get remodeled in just a few seconds (Lacy et al., 2019; Watanabe and Mitchison, 2002). This suggests the existence of intracellular mechanisms for accelerating phosphate release (Blanchoin and Pollard, 1999) and/or directly disassembling ADP-P_i_ filaments, thus bypassing the slow P_i_ release step.

Indeed, recent findings demonstrate that mouse twinfilin accelerates the depolymerization of newly-assembled (ADP-P_i_) actin filaments, specifically at the filament barbed ends (Shekhar et al., 2021). However, the question of whether cofilin together with one of its main co-factors, cyclase-associated protein, can promote pointed-end depolymerization of ADP-P_i_ filaments remains unresolved.

Cofilin has long been considered the central player in actin disassembly (Carlier et al., 2015; Goode et al., 2023; Lappalainen and Drubin, 1997). It binds filaments along their sides, accelerates phosphate release by up to 10-fold (Blanchoin and Pollard, 1999), severs filaments (Maciver et al., 1991) and promotes depolymerization at both filament ends (Shekhar and Carlier, 2017; Wioland et al., 2017). Importantly, saturation of filament sides by cofilin accelerates pointed-end depolymerization and decelerates barbed-end depolymerization (Wioland et al., 2017). The barbed-end targeting of undecorated filaments by cofilin is responsible for accelerating barbed-end depolymerization (Wioland et al., 2017). While both budding yeast (*S. cerevisiae*) and fission yeast (*S. pombe*) only express a single cofilin, mammals express three different isoforms. These include the ubiquitously expressed cofilin-1, actin depolymerizing factor (ADF), which is primarily found in neuronal, epithelial, and endothelial tissues, and muscle-specific cofilin-2 (together known as ADF/cofilin) (Vartiainen et al., 2002). All three mammalian cofilins sever and modestly accelerate depolymerization of ADP filaments (Chin et al., 2016; Shekhar and Carlier, 2017; Wioland et al., 2017).

Cofilin is assisted in actin disassembly by its cofactors such as Coronin, Aip1 and cyclase-associated protein (CAP) (Goode et al., 2023). CAP is a multifunctional actin-binding protein conserved across animals, plants, and fungi (Ono, 2013). CAP monomers consist of two halves; each half is capable of functioning autonomously (Chaudhry et al., 2014). The C-terminal half (or C-CAP) consists of actin-binding WH2 and β-sheet/CARP domains as well as poly-proline stretches capable of binding profilin. C-CAP promotes the dissociation of cofilin from cofilin-bound ADP-actin subunits released from filament pointed-ends and stimulates nucleated exchange (from ADP to ATP) to recycle actin monomers for new rounds of polymerization (Balcer et al., 2003; Chaudhry et al., 2010; Jansen et al., 2014; Kotila et al., 2018; Makkonen et al., 2013; Mattila et al., 2004; Moriyama and Yahara, 2002). C-CAP also enhances barbed-end depolymerization (Alimov et al., 2023; Towsif and Shekhar, 2023). The N-terminal half of CAP (referred to as N-CAP) on its own modestly promotes pointed-end depolymerization of ADP actin filaments (Moriyama and Yahara, 2002; Shekhar et al., 2019). N-CAP further synergizes with cofilin to accelerate pointed-end depolymerization of ADP filaments by up to 300-fold (Kotila et al., 2019; Shekhar et al., 2019). The role of N-CAP in promoting cofilin-mediated severing is a topic of ongoing debate. While two earlier studies employing conventional TIRF microscopy indicated that N-CAP and cofilin could together enhance severing by up to 8-fold (Chaudhry et al., 2013; Jansen et al., 2014), a more recent mf-TIRF study failed to observe any increase in cofilin-mediated severing induced by N-CAP (Kotila et al., 2019). Notwithstanding these disagreements on severing, depolymerization of ADP actin filaments by cofilin and N-CAP is relatively well-understood. Whether these proteins, either individually or in tandem, can also depolymerize ADP-P_i_ filaments still remains unresolved.

Here we show that mammalian cofilin promotes pointed-end depolymerization of ADP-P_i_ filaments. Importantly, we find that this behavior is preserved across all three human isoforms of cofilin and requires cofilin’s ability to interact with sides of actin filaments. We also show that synergistic depolymerization by N-CAP and cofilin persists for ADP-P_i_ filaments. The absolute rate of depolymerization for ADP-P_i_ filaments however is much slower than previously reported rates for ADP filaments (Kotila et al., 2019; Shekhar et al., 2019). Notably, we find that neither protein on its own nor together promotes severing of ADP-P_i_ filaments.

**Fig. 1:**
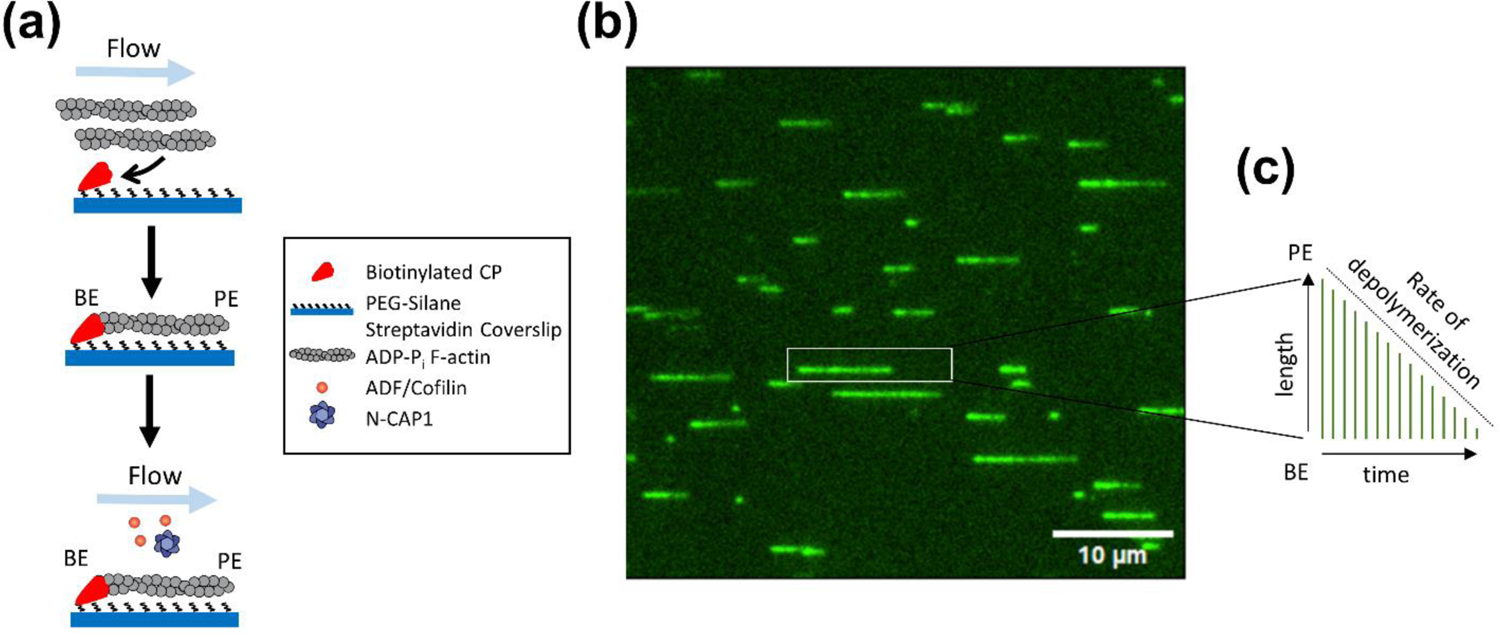
Microfluidics-assisted TIRF microscopy (mf-TIRF) for studying pointed-end depolymerization of ADP-P_i_ actin filaments (a) Schematic of the experimental strategy. Pre-formed Alexa-488 labeled ADP-P_i_ actin filaments were captured by coverslip-anchored biotinylated capping protein in the mf-TIRF chamber. These filaments were exposed to a flow containing cofilin isoforms and/or N-CAP1 (or control buffer) in presence of 50 mM P_i_. Depolymerization of free pointed ends was monitored. BE, barbed end; PE, pointed end. **(b)** Representative field of view showing CP-anchored actin filaments aligned along the flow. **(c)** The methodology used for determining the rate of pointed-end depolymerization from kymograph analysis.

## Results and Discussion

### Cofilin accelerates pointed-end depolymerization of ADP-P_i_ filaments

To determine the effects of human cofilin-1 on the pointed-end dynamics of newly assembled actin filaments, we compared the time-dependent length changes of CP-capped ADP-Pi actin filaments in the presence and absence of cofilin-1. Preformed fluorescently-labeled actin filaments were introduced into the mf-TIRF microfluidic chamber and captured at their barbed ends by capping protein (CP) bound to the glass coverslip (Fig. 1a). Anchoring filaments via CP specifically ensured that any depolymerization-induced filament length changes occurred only due to dynamics at free pointed ends. The mf-TIRF approach facilitated the simultaneous recording of pointed-end dynamics of a large number of filaments with high temporal and spatial resolution (Fig. 1b)(Shekhar and Carlier, 2016). To maintain actin subunits in the filaments in the ADP-P_i_ state, all reactions were supplemented with 50 mM P_i_ (Carlier and Pantaloni, 1988; Shekhar et al., 2021). Time-lapse images were recorded and the rate of pointed-end depolymerization was determined using kymograph analysis (Fig. 1c). In control reactions, we found the rate of pointed-end depolymerization to be undetectably slow (Fig. 2a). In the presence of cofilin-1, however, an increase in the rate of depolymerization was observed (Fig. 2b). The rate increased linearly with cofilin-1 concentration (Fig. 2c). A similar concentration-dependent behavior was previously reported for pointed-end depolymerization of ADP-actin filaments by actin depolymerizing factor (ADF) (Shekhar and Carlier, 2017). Importantly, the depolymerization rate of ADP filaments saturated upon complete decoration of the filament by ADF/cofilin (Wioland et al., 2017). We did not detect a saturation of the depolymerization rate even at high cofilin-1 concentrations. At 20 µM cofilin-1, the highest concentration we were able to test, we measured a depolymerization rate of 0.55 ± 0.11 su/s (Fig. 2c), very close to the rate reported for cofilin-1-saturated ADP-actin filaments at the same pH of 7.4 (Wioland et al., 2017; Wioland et al., 2019).

**Fig. 2:**
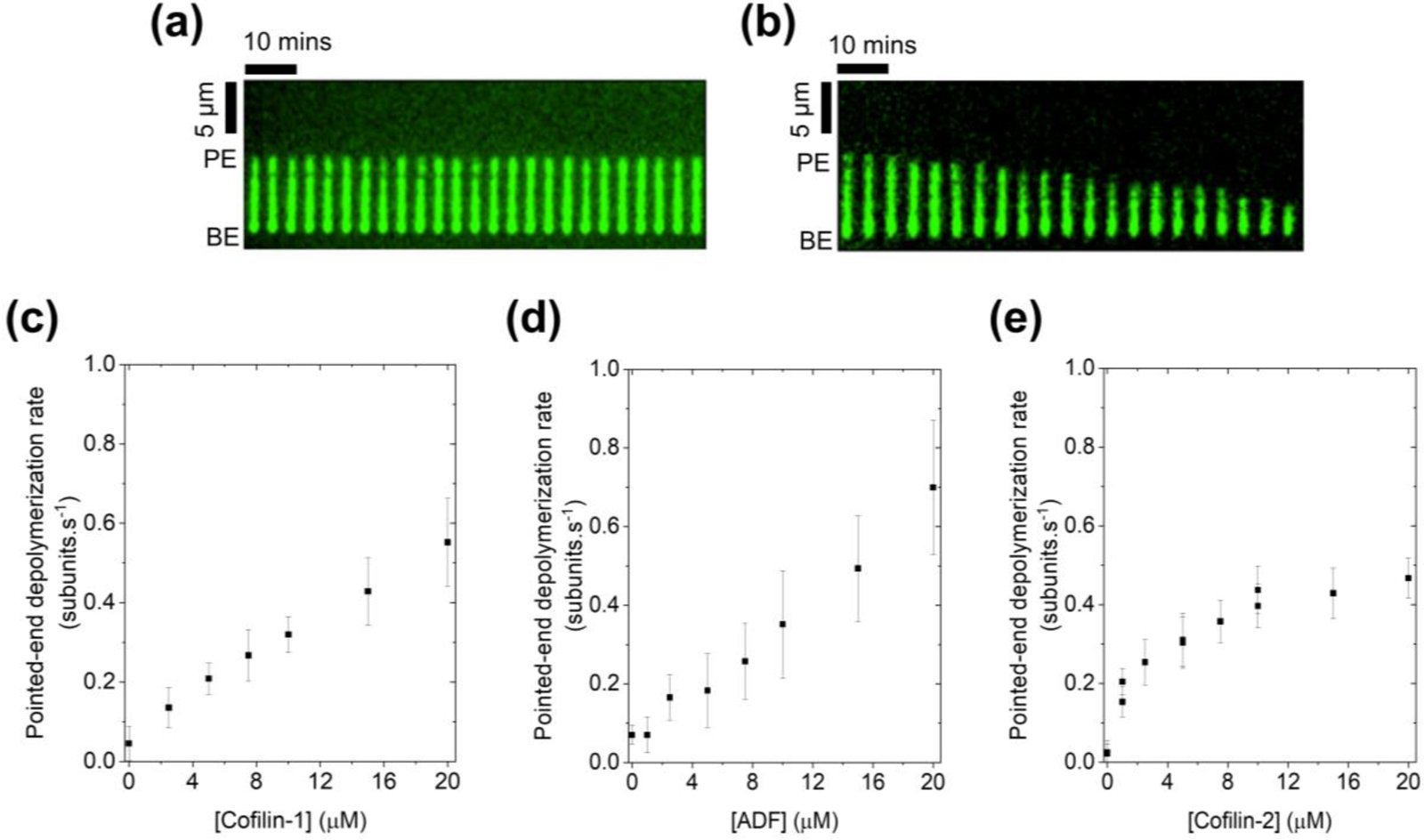
Pointed-end depolymerization of ADP-P_i_ actin filaments is conserved for all mammalian cofilin isoforms. (a) Kymograph of an Alexa-488 labeled ADP-P_i_ actin filament depolymerizing in TIRF buffer containing 50 mM P_i_ **(b)** Same as (a) but with 20 μM cofilin-1. **(c)** Rates (± sd) of pointed-end depolymerization in the presence of varying concentrations of cofilin-1. **(d)** Rates (± sd) of pointed-end depolymerization in the presence of varying concentrations of actin depolymerizing factor. **(e)** Rates (± sd) of pointed-end depolymerization in the presence of varying concentrations of cofilin-2. N = 20 – 68 filaments were analyzed for each concentration.

Mammals express two additional cofilin isoforms, namely ADF and cofilin-2. We therefore sought to investigate whether these effects were conserved across isoforms. Our examination of ADF revealed a concentration-dependent trend similar to cofilin-1 (Fig. 2d). In contrast, the depolymerization rate for cofilin-2 appeared to saturate at high concentrations (Fig. 2e), consistent with a previous study demonstrating that cofilin-2 exhibits a higher affinity for ATP-G-actin and more efficiently accelerates the disassembly of ADP⋅BeFx-actin filaments than cofilin-1 (Kremneva et al., 2014). This investigation from the Lappalainen lab also identified specific residues in cofilin-2 crucial for binding ADP-P_i_-actin filaments and ATP-actin monomers. Thus, mirroring previous observations on ADP actin filaments (Wioland et al., 2017), while all three cofilin isoforms display the capacity to induce pointed-end depolymerization of ADP-P_i_ filaments, cofilin-2 may have evolved to effectively disassemble actin filaments in muscle sarcomeres, potentially containing newly assembled actin subunits at their pointed ends (Littlefield et al., 2001).

### Pointed-end depolymerization of ADP-P_i_ filaments requires binding of cofilin to filament sides

We then sought to determine whether cofilin-mediated depolymerization of ADP-P_i_ filaments requires binding of cofilin to filament sides, similar to ADP actin filaments. However, due to the high concentration of cofilin needed for detectable pointed-end depolymerization of ADP-P_i_ filaments, direct visualization of fluorescently-labeled cofilin interacting with actin filaments was not feasible. Consequently, we conducted co-sedimentation experiments. Pre-assembled actin filaments were incubated with various cofilin concentrations in the absence or presence of 50 mM Pi. Consistent with earlier findings (Carlier et al., 1997), cofilin readily pelleted with F-actin in the absence of Pi (Fig. 3a,b). As the total cofilin concentration approached the F-actin concentration, amount of cofilin found in the pellet increased until no further increase in pelleted cofilin was seen, presumably due to complete saturation of F-actin by cofilin. When we repeated these experiments in presence of 50 mM P_i_, we found that at comparable cofilin concentrations, the amount of cofilin found in the pellet was consistently less for ADP-P_i_ filaments than what was observed for ADP filaments. We note that we didn’t observe saturation (i.e. complete decoration) of ADP-P_i_ by cofilin up to an incubation concentration of 20 µM cofilin, indicating that only a fraction of ADP-P_i_ F-actin subunits were bound to cofilin even at high cofilin concentrations Our observations align with the previously reported preferential binding of cofilin to ADP actin over ADP-P_i_ actin (Blanchoin and Pollard, 1999).

**Fig. 3:**
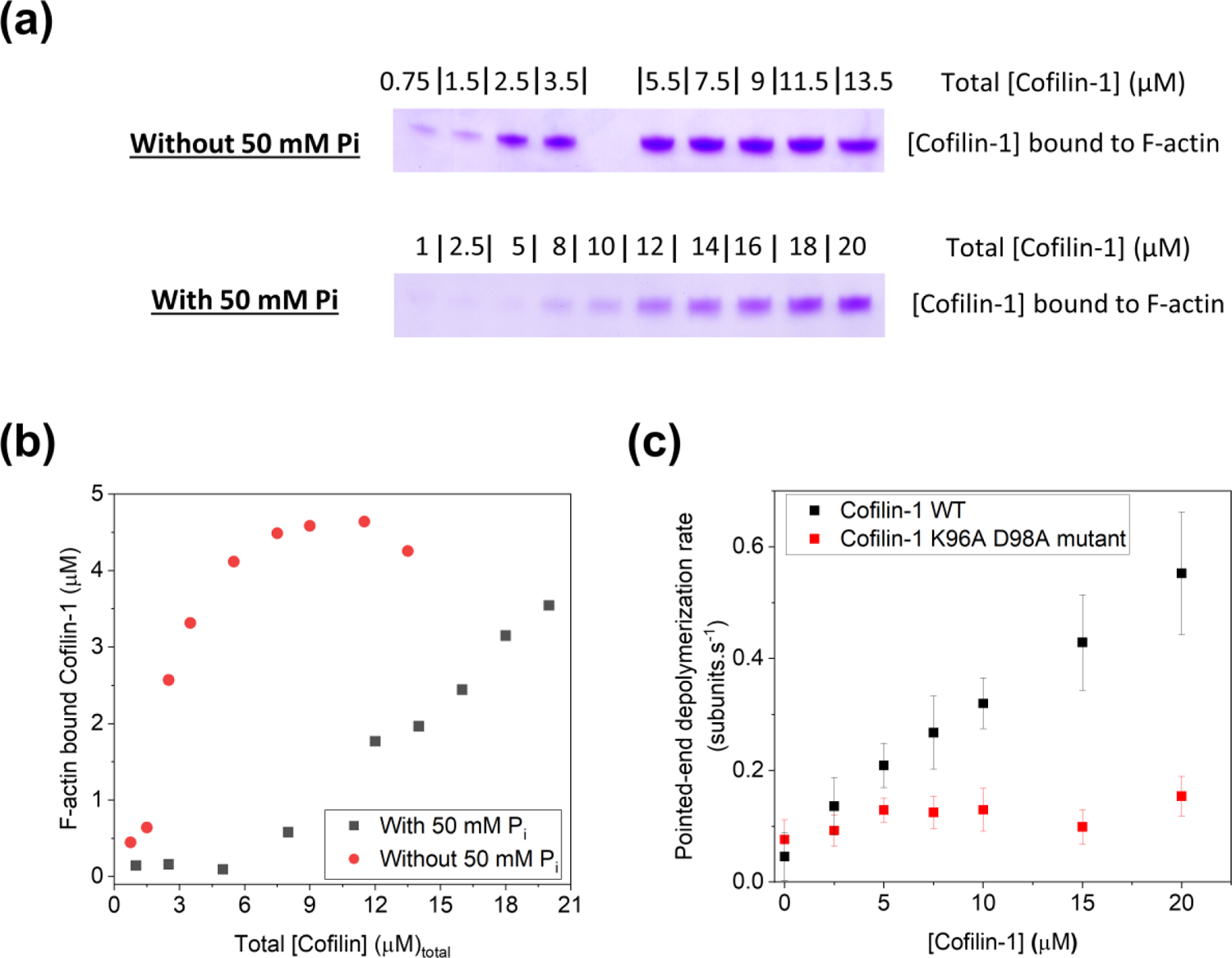
Binding of cofilin to sides of actin filaments is required for cofilin-mediated depolymerization of ADP-P_i_ (a) SDS-PAGE of cofilin-1 in the pellet of sedimented samples containing 5 µM F-actin and varying amounts of cofilin (total incubation concentration indicated). Left: in absence of 50 mM Pi; Right: in presence of 50 mM Pi. **(b)** Concentration of cofilin-1 in the pellet after incubation with 5 µM F-actin in absence (black symbols) or presence of 50 mM Pi (red symbols) **(c)** Rates (± sd) of pointed-end depolymerization in the presence of varying concentrations of wild-type (black) and K96A D98A mutant cofilin-1 (red). N = 25– 28 filaments were analyzed for each concentration. Note that the data for wild-type cofilin is the same as shown in Fig. 2c.

To further validate our hypothesis that side binding of cofilin was critical for pointed-end depolymerization, we expressed and purified a cofilin mutant (K96A D98A) with a weakened affinity for filament sides owing to point-mutations in its F-actin binding region (Lappalainen et al., 1997; Wioland et al., 2017). When the actin filaments were exposed to the mutant cofilin, a significant decrease in the depolymerization rate was observed compared to the wild-type protein (Fig. 3c). This observation lends support to the essential role of cofilin’s F-actin binding in inducing depolymerization of ADP-P_i_ filaments. Nevertheless, our experiments do not definitively exclude the possibility although cofilin can bind sides of actin subunits in the bulk of the filament, depolymerization might result solely from cofilin’s direct targeting of terminal actin subunit(s) at filament pointed ends.

### Cofilin-1 does not promote severing of ADP-P_i_ actin filaments

Binding of cofilin alters the conformation and mechanical properties of actin filaments (Elam et al., 2013; Galkin et al., 2011; Huehn et al., 2018). Discontinuities between cofilin-decorated and ‘bare’ regions on filaments cause their severing (Suarez et al., 2011). Based on our observations that cofilin can indeed bind sides of ADP-P_i_ filaments, albeit with less efficiency, we anticipated that the partially decorated ADP-P_i_ filaments would display elevated rates of cofilin-mediated severing. However, contrary to our expectations and in agreement with previous studies on Acanthamoeba actophorin (Maciver et al., 1991), no noticeable severing was observed within the tested range of cofilin concentrations (0 to 20 µM). These findings suggest that although cofilin can bind to the sides of both ADP and ADP-P_i_ actin filaments, its ability to induce severing is specifically confined to aged actin filaments.

The process of cofilin-mediated severing is proposed to involve multiple steps—cofilin initially binds to F-actin subunits, followed by an isomerization step in the cofilin-bound subunit that facilitates cooperative binding of cofilin on neighboring actin subunits (De La Cruz and Sept, 2010; Hayakawa et al., 2014). Cooperative binding of cofilin is expected to result in the formation of cofilin-decorated and bare regions on an actin filament, leading to severing between these regions (Suarez et al., 2011). We speculate that the cooperative binding of cofilin might be specifically limited to ADP filaments and not ADP-P_i_ filaments. Consequently, although cofilin may transiently bind to ADP-P_i_ F-actin subunits, the lack of cooperativity may hinder the formation of long stretches of cofilin-coated interspersed with cofilin-free bare regions. Further insight into this phenomenon would require future cryo-EM studies to uncover the structural changes induced by cofilin on ADP-P_i_ filaments. This would help resolve why cofilin can depolymerize both ADP-P_i_ and ADP filaments but can only sever the latter.

### Synergistic depolymerization by N-CAP and cofilin persists for ADP-P_i_ filaments

Cyclase-associated protein (CAP) enhances pointed-end depolymerization of ADP actin filaments through the activity of its N-terminal half (or N-CAP) (Kotila et al., 2019; Moriyama and Yahara, 2002; Shekhar et al., 2019). We therefore first asked if N-CAP was also capable of similarly enhancing depolymerization of ADP-P_i_ pointed ends. When we exposed ADP-P_i_ filament pointed ends to N-CAP1, we observed no significant enhancement of pointed-end depolymerization. Consistent with previous findings regarding ADP filaments (Jansen et al., 2014), N-CAP1 also did not induce any measurable severing of ADP-P_i_ filaments.

N-CAP and cofilin have previously been shown to synergize in accelerating pointed-end depolymerization of ADP-actin filaments by up to 300-fold (Kotila et al., 2019; Shekhar et al., 2019). To determine if N-CAP-cofilin synergy might extend to ADP-P_i_ filaments, we first systematically tuned the concentration of cofilin-1 in the range of 0 to 10 µM while keeping N-CAP1 concentration fixed at 5 µM (monomers). The recorded depolymerization rates were then compared with those observed for cofilin-1 alone. In both the presence and absence of N-CAP1, we observed a concentration-dependent increase in the rate of pointed-end depolymerization (Fig. 4a). Across the entire concentration range, the addition of N-CAP1 resulted in a 2-fold increase in pointed-end depolymerization compared to reactions with cofilin-1 alone. Further analysis revealed that at a fixed concentration of cofilin-1 (5 µM), increasing N-CAP1 concentration led to an increase in pointed-end depolymerization (Fig. 4b). To delve deeper into the mechanism of cofilin: N-CAP1 synergy, we repeated these experiments while maintaining cofilin-1 concentration fixed at 10 µM or 20 µM. Similar to the experiment with 5 µM cofilin-1, N-CAP1 once again increased depolymerization in a concentration-dependent fashion (Fig. 4b). The maximal rate of depolymerization increased with rising concentrations of cofilin (Fig. 4c). Notably, the depolymerization effect for all three cofilin concentrations (5, 10, and 20 µM) saturated at approximately 4 µM N-CAP1. While the cytosolic concentration of CAP in mammalian cells is currently unknown, the concentration of N-CAP1 required for maximal depolymerization in each condition closely approximates the cellular concentration of CAP in *S. cerevisiae* (3 μM monomers) (Johnston et al., 2015).

Considering our co-sedimentation experiments alongside our kinetics data, we propose that, at sub-saturating concentrations of cofilin and saturating concentrations of N-CAP1, the net depolymerization rate is defined by the probability of terminal actin subunits being bound to both cofilin and N-CAP1. As the cofilin concentration increases, the probability of the terminal actin subunits being bound to cofilin is expected to increase. These cofilin-bound subunits, in turn, serve as substrates for the accelerated depolymerization action of N-CAP1, leading to faster overall depolymerization. Consistently, the filaments displayed lower maximal depolymerization when wild-type cofilin was replaced by a cofilin mutant (both in presence of N-CAP) with reduced affinity for actin subunit’s F-site (Fig. 4d). We note that our results do not distinguish how terminal actin gets bound to cofilin – whether due to decoration of filament sides by cofilin or due to specific targeting terminal actin subunits by cofilin.

In conjunction with previous studies, our data suggest that cofilin and N-CAP1 work synergistically to promote pointed-end depolymerization, irrespective of nucleotide state. Importantly, we note that the maximal rates achieved at comparable concentrations of mammalian cofilin and N-CAP1 (alone and together) are significantly lower for ADP-P_i_ filaments compared to ADP actin filaments. Additionally, unlike observations on ADP actin filaments (Chaudhry et al., 2013; Jansen et al., 2014), we did not observe any increase in severing of ADP-P_i_ filaments with the simultaneous presence of cofilin and N-CAP1.

**Fig. 4:**
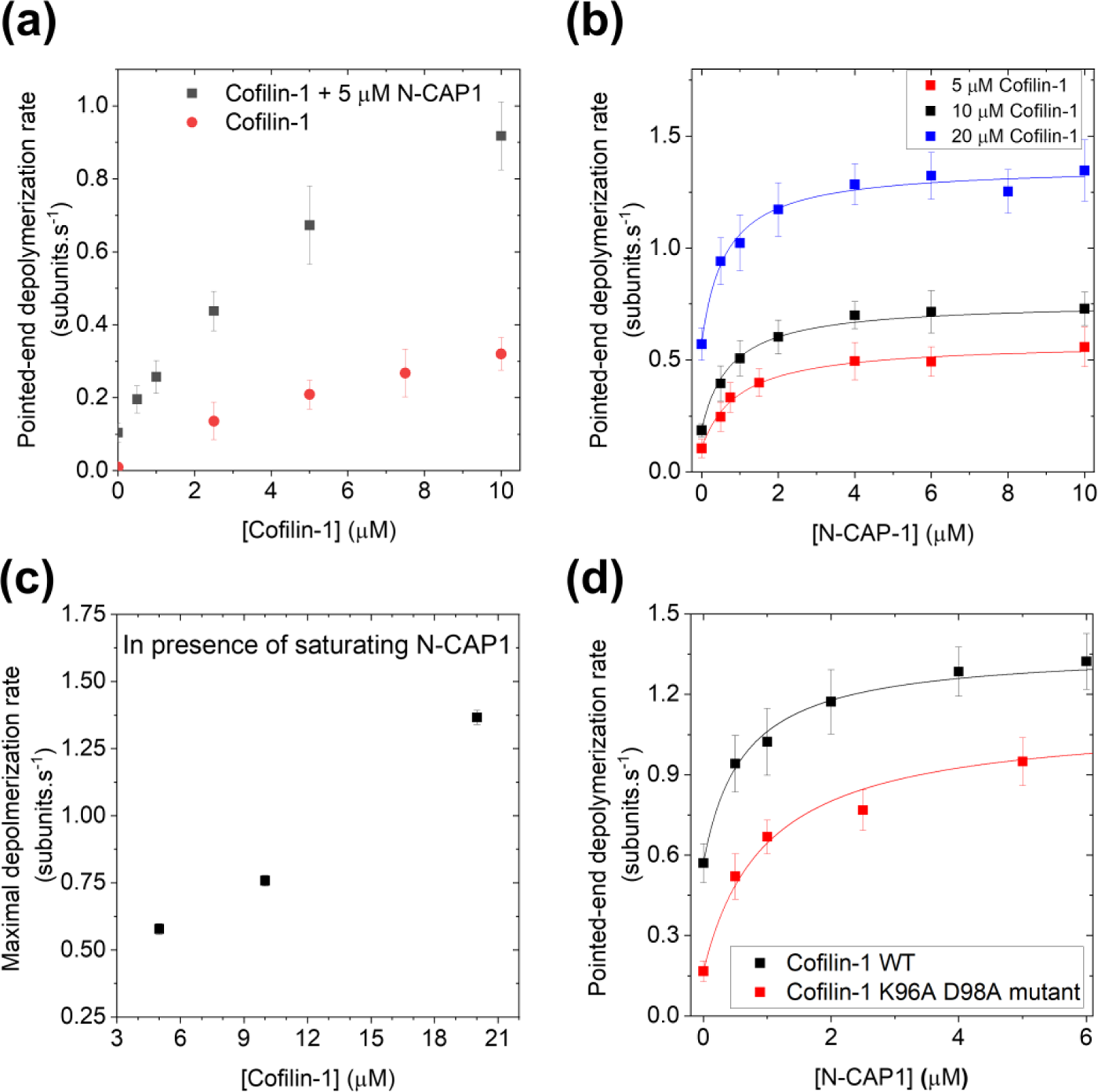
Cofilin-1 and N-CAP1 synergize to accelerate ADP-P_i_ pointed-end depolymerization (a) Rates (± sd) of pointed-end depolymerization as a function of cofilin-1 concentration in the presence (black symbols) or absence (red symbols) of 5 µM N-CAP1. All stated N-CAP1 concentrations are for monomers. **(b)** Rates (± sd) of pointed-end depolymerization as a function of N-CAP1 concentration in the presence of 5 µM cofilin-1 (red symbols), 10 µM cofilin-1 (black symbols) and 20 µM cofilin-1 (blue symbols). *N* = 20 - 26 filaments analyzed per condition. Data were fit to a saturation binding model (see methods). K_d_ determined from the fits were as follows – 0.62 ± 0.11 µM (for 20 µM Cofilin-1), 0.75 ± 0.13 µM (for 10 µM Cofilin-1) and 0.96 ± 0.13 µM (for 5 µM Cofilin-1) **(c)** Maximal rate of pointed-end depolymerization as a function of cofilin-1 concentration in presence of saturating N-CAP1 as determined from data shown in (c). **(d)** Rates (± sd) of pointed-end depolymerization as a function of N-CAP1 concentration in the presence of 20 µM cofilin-1 wild type (black) or K96A D98A mutant cofilin-1. N = 23– 25 filaments were analyzed for each concentration. Note that the data for wild-type cofilin-1 and N-CAP1 condition is the same as shown in Fig. 4c. Data were fit (lines) to a saturation binding model (see methods). K_d_ determined from the fits were as follows – 0.62 ± 0.11 µM (for 20 µM wild-type Cofilin-1) and 0.97 ± 0.16 (for 20 µM mutant Cofilin-1).

## Conclusions

By integrating our findings with previous research, we propose a working model illustrating the nucleotide-specific disassembly of actin filaments through severing and enhanced depolymerization by cofilin and CAP (Fig. 5) at the two ends of actin filaments. Cofilin can decorate the sides of both aged and newly-assembled actin filaments. Our data, in agreement with the earlier study by Blanchoin and Pollard, suggest that the extent of filament decoration by cofilin strongly relies on the nucleotide state (Blanchoin and Pollard, 1999). Consequently, ADP actin filaments exhibit a lower threshold of cofilin concentration required for complete decoration compared to ADP-P_i_ actin filaments. Similarly, the depolymerizing effects of CAP in presence of cofilin, are more pronounced in aged ADP actin filaments than in newly-assembled ADP-P_i_ filaments.

**Fig. 5:**
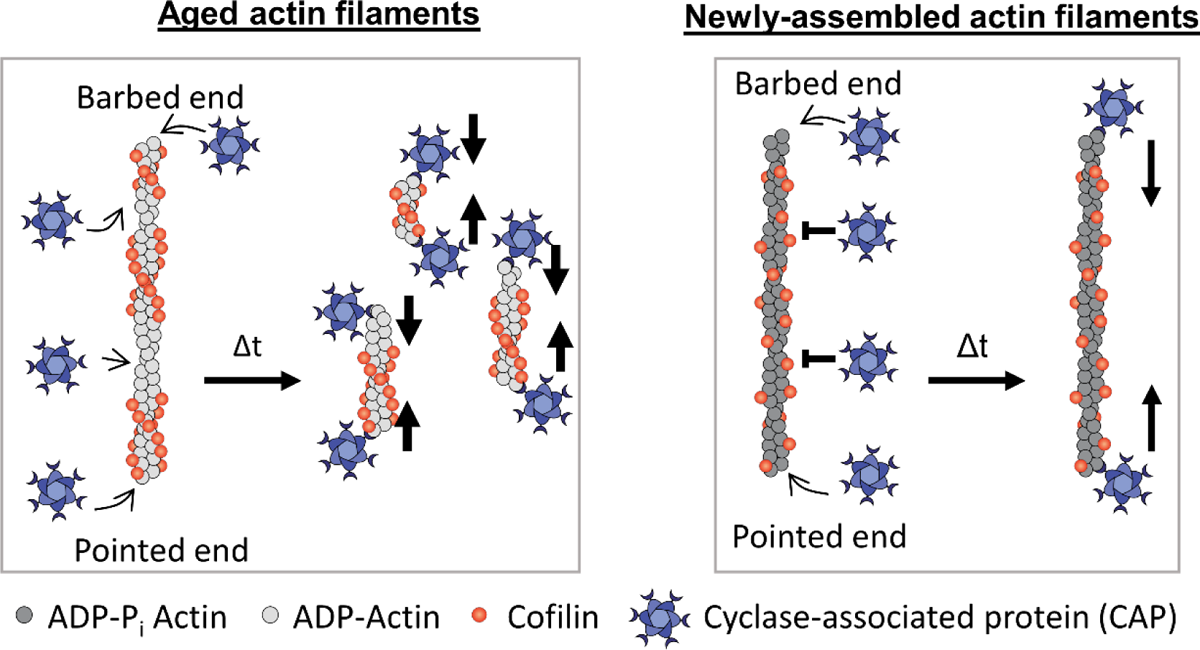
Working model of synergistic depolymerization by ADF/Cofilin and CAP as a function of filament age. Synergistic depolymerization by simultaneous action of cofilin and CAP persists for ADP (left) and ADP-P_i_ filaments (right). Cofilin (alone and together with CAP) efficiently severs ADP-but not ADP-P_i_ filaments. Severing occurs at boundaries between cofilin-decorated and bare actin filament (Suarez et al., 2011). Since cooperative binding of cofilin has been thought to underlie formation of cofilin-decorated and bare regions, we postulate that cofilin’s binding to ADP-P_i_ filaments might not be cooperative which results in cofilin randomly distributed along the filament length rather than in stretches responsible for formation of cofilin-bound and bare regions (De La Cruz and Sept, 2010). In both cases, CAP is able to depolymerize barbed ends (Alimov et al., 2023; Towsif and Shekhar, 2023).

While our current study demonstrates cofilin-mediated depolymerization occurring on both aged and newly-assembled actin filaments, filament severing appears to be specific to aged actin filaments. Other factors, including coronin and Aip1, have been reported to enhance cofilin’s binding to aged actin filaments, promoting severing in cofilin-saturated filaments that would otherwise not sever (Gressin et al., 2015; Jansen et al., 2015; Nadkarni and Brieher, 2014; Okada et al., 1999; Okada et al., 2006; Rodal et al., 1999). Future investigations are necessary to understand how the collaborative disassembly of actin filaments by cofilin, coronin, and Aip1 may be influenced by the age of actin filaments (Jansen et al., 2015).

In addition to its synergistic depolymerization with cofilin, CAP acts in concert with another member of the ADF/cofilin family protein twinfilin to depolymerize barbed and pointed ends in a species-specific manner. *S. cerevisiae* twinfilin collaborates with Srv2 (CAP homolog in *S. cerevisiae*) to accelerate depolymerization of both barbed and pointed ends by 3-fold and 17-fold respectively (Johnston et al., 2015). The collaboration between mouse twinfilin and CAP in accelerating barbed-end depolymerization, however, remains a subject of controversy. While an earlier study presented evidence supporting their synergistic action (Hilton et al., 2018), a more recent study failed to find evidence of N-CAP accelerating filament barbed-end depolymerization by twinfilin (Hakala et al., 2021). In the future, it will be crucial to investigate whether, akin to cofilin-CAP synergy, twinfilin-CAP synergy can also contribute to the depolymerization of newly-assembled actin filaments.

Where in a cell might depolymerization of ADP-P_i_ pointed ends be relevant? Firstly, in structures like lamellipodia where actin networks turn over much faster than the time required for P_i_ release (Watanabe and Mitchison, 2002). Secondly, in processes where ATP-actin subunits are being added directly at filament pointed-ends. Until recently, pointed-ends *in vivo* were only thought to be capable of depolymerization. Recent discover of processive pointed-end polymerization of ATP-actin subunits at filament pointed ends by the bacterial effector protein VopF have opened the possibility of similar eukaryotic mechanisms (Kudryashova et al., 2022). Polymerization at pointed-ends would generate actin filaments rich in ADP-P_i_ actin subunits, and the CAP-Cofilin synergistic depolymerization mechanism described here could contribute to their depolymerization. In summary, the mechanisms outlined here introduce the exciting prospect of rapidly disassembling newly-assembled actin networks without undergoing the aging process.

## Materials and methods

### Purification and labeling of actin

Rabbit skeletal muscle actin was purified from acetone powder generated from frozen ground hind leg muscle tissue of young rabbits (PelFreez, USA). Lyophilized acetone powder stored at −80°C was mechanically sheared in a coffee grinder, resuspended in G-buffer (5 mM Tris-HCl pH 7.8, 0.5 mM Dithiothreitol (DTT), 0.2 mM ATP and 0.1 mM CaCl_2_), and cleared by centrifugation for 20 min at 50,000 × *g*. Supernatant was collected and further filtered with Whatman paper. Actin was then polymerized overnight at 4°C, slowly stirring, by the addition of 2 mM MgCl_2_ and 50 mM NaCl to the filtrate. The next morning, NaCl powder was added to a final concentration of 0.6 M and stirring was continued for another 30 min at 4°C. Then, F-actin was pelleted by centrifugation for 150 min at 280,000 × *g*, the pellet was solubilized by dounce homogenization and dialyzed against G-buffer for 48 h at 4°C. Monomeric actin was then precleared at 435,000 × *g* and loaded onto a Sephacryl S-200 16/60 gel-filtration column (Cytiva, USA) equilibrated in G-Buffer. Fractions containing actin were stored at 4°C.

To fluorescently label actin, G-actin was polymerized by dialyzing overnight against modified F-buffer (20 mM PIPES pH 6.9, 0.2 mM CaCl_2,_ 0.2 mM ATP, 100 mM KCl) (Shekhar, 2017). F-actin was incubated for 2 h at room temperature with a 5-fold molar excess of Alexa-488 NHS ester dye (Thermo Fisher Scientific, USA). F-actin was then pelleted by centrifugation at 450,000 × *g* for 40 min at room temperature, and the pellet was resuspended in G-buffer, homogenized with a dounce and incubated on ice for 2 h to depolymerize the filaments. The monomeric actin was then re-polymerized on ice for 1 h by addition of 100 mM KCl and 1 mM MgCl_2_. F-actin was once again pelleted by centrifugation for 40 min at 450,000 × *g* at 4°C. The pellet was homogenized with a dounce and dialyzed overnight at 4°C against 1 L of G-buffer. The solution was precleared by centrifugation at 450,000 × *g* for 40 min at 4°C. The supernatant was collected, and the concentration and labeling efficiency of actin was determined.

### Purification of profilin

Human profilin-1 was expressed in *E. coli* strain BL21 (pRare) to log phase in LB broth at 37°C and induced with 1 mM IPTG for 3 h at 37°C. Cells were then harvested by centrifugation at 15,000 × *g* at 4°C and stored at -80°C. For purification, pellets were thawed and resuspended in 30 mL lysis buffer (50 mM Tris-HCl pH 8, 1 mM DTT, 1 mM PMSF protease inhibitors (0.5 μM each of pepstatin A, antipain, leupeptin, aprotinin, and chymostatin)) was added, and the solution was sonicated on ice by a tip sonicator. The lysate was centrifuged for 45 min at 120,000 × *g* at 4°C. The supernatant was then passed over 20 ml of Poly-L-proline conjugated beads in a disposable column (Bio-Rad, USA). The beads were first washed at room temperature in wash buffer (10 mM Tris pH 8, 150 mM NaCl, 1 mM EDTA and 1 mM DTT) and then washed again with 2 column volumes of 10 mM Tris pH 8, 150 mM NaCl, 1 mM EDTA, 1 mM DTT and 3 M urea. Protein was then eluted with 5 column volumes of 10 mM Tris pH 8, 150 mM NaCl, 1 mM EDTA, 1 mM DTT and 8 M urea. Pooled and concentrated fractions were then dialyzed in 4 L of 2 mM Tris pH 8, 0.2 mM EGTA, 1 mM DTT, and 0.01% NaN_3_ (dialysis buffer) for 4 h at 4°C. The dialysis buffer was replaced with fresh 4 L buffer and the dialysis was continued overnight at 4°C. The protein was centrifuged for 45 min at 450,000 × *g* at 4°C, concentrated, aliquoted, flash frozen in liquid N_2_ and stored at -80°C.

### Purification of wild-type and mutant ADF/cofilins

Human cofilin-1, cofilin-2,actin depolymerizing factor (ADF) and K96A D98A mutant cofilin-1 were expressed in *E.coli* BL21 DE3 cells. Cells were grown in Terrific Broth to log phase at 37⁰C, and then expression was induced overnight at 18⁰C by addition of 1 mM IPTG. Cells were collected by centrifugation and pellets were stored at -80⁰C. Frozen pellets were thawed and resuspended in lysis buffer (20 mM Tris pH 8.0, 50 mM NaCl, 1 mM DTT, and protease inhibitors (0.5 µM each of pepstatin A, antipain, leupeptin, aprotinin, and chymostatin). Cells were lysed with a tip sonicator while being kept on ice. The cell lysate was centrifuged at 150,000 × *g* for 30 min at 4⁰C. The supernatant was loaded on a 5 ml HiTrap Q HP column (GE Healthcare, Pittsburgh, PA), and the flow-through was collected and dialyzed against 20 mM HEPES pH 6.8, 25 mM NaCl, and 1 mM DTT. The dialyzed solution was then loaded on a 5 ml HiTrap SP FF column (GE Healthcare, Pittsburgh, PA) and eluted using a linear gradient of NaCl (20-500 mM). Fractions containing protein were concentrated, dialyzed against 20 mM Tris pH 8.0, 50 mM KCl, and 1 mM DTT, flash frozen in liquid N_2_ and stored at -80⁰C.

### Purification and biotinylation of SNAP-CP

SNAP-tagged capping protein (Bombardier et al., 2015) was expressed in *E. coli* BL21 DE3 by growing cells to log phase at 37°C in TB medium, then inducing expression using 1 mM IPTG at 18°C overnight. Cells were harvested by centrifugation and pellets were stored at −80°C. Frozen pellets were resuspended in lysis buffer (20 mM NaPO_4_ pH 7.8, 300 mM NaCl, 1 mM DTT, 15 mM imidazole, 1 mM PMSF) supplemented with a protease inhibitor cocktail (0.5 µM each of pepstatin A, antipain, leupeptin, aprotinin, and chymostatin). Cells were lysed by sonication with a tip sonicator while keeping the tubes on ice. The lysate was cleared by centrifugation at 150,000 x *g* for 30 min at 4°C. The supernatant was then flowed through a HisTrap column connected to a Fast Protein Liquid Chromatography (FPLC) system. The column with the bound protein was first extensively washed with the washing buffer (20 mM NaPO_4_ pH 7.8, 300 mM NaCl, 1 mM DTT and 15 mM imidazole) to remove non-specifically bound proteins. SNAP-CP was then eluted with a linear zero to 250 mM imidazole gradient in 20 mM NaPO_4_ pH7.8, 300 mM NaCl, and 1 mM DTT. The eluted protein was concentrated and labelled with Benzylguanine-Biotin (New England Labs) according to the manufacturer’s instructions. Free biotin was removed using size-exclusion chromatography by loading the labelled protein on a Superose 6 gel filtration column (GE Healthcare, Pittsburgh, PA) eluted with 20 mM HEPES pH 7.5, 150 mM KCl, 0.5 mM DTT. Fractions containing the protein were combined, aliquoted, snap frozen in liquid N_2_ and stored at -80 °C.

### Purification of N-CAP1

His-tagged N-terminal fragment mouse N-CAP1 (Jansen et al., 2014) was expressed in *E. coli* BL21 DE3 by growing cells to log phase at 37°C in TB medium. Cells were induced with 1 mM IPTG at 18°C overnight. Cells were harvested by centrifugation and pellets were stored at −80°C. Frozen pellets were resuspended in 50 mM NaPO_4_ pH 8.0, 1 mM PMSF, 1 mM DTT, 20 mM imidazole, 300 mM NaCl and protease inhibitors as described above. Cells were lysed by sonication with a tip sonicator while keeping the tubes on ice. The lysate was cleared by centrifugation at 150,000 x g for 30 min at 4°C. The lysate was loaded on a 1 ml HisTrap HP column (GE Healthcare, Pittsburgh, PA) and non-specifically bound proteins were removed by washing the column with 20 mM NaPO_4_ pH 8.0, 50 mM imidazole, 300 mM NaCl and 1 mM DTT. The bound protein was then eluted using a linear gradient of 50 – 250 mM imidazole in the same buffer. Fractions containing the protein were concentrated and dialyzed into 10 mM imidazole pH 8.0, 150 mM NaCl and 1 mM DTT. The protein was then aliquoted, snap frozen in liquid N_2_ and stored at -80 °C.

### Microfluidics-assisted TIRF (mf-TIRF) microscopy

Actin filaments were first assembled in microfluidics-assisted TIRF (mf-TIRF) flow cells (Jegou et al., 2011; Shekhar, 2017). For all experiments, coverslips were first cleaned by sonication in Micro90 detergent for 20 min, followed by successive 20 min sonications in 1 M KOH, 1 M HCl and 200 proof ethanol for 20 min each. Washed coverslips were then stored in fresh 200 proof ethanol. Coverslips were then washed extensively with H_2_O and dried in an N_2_ stream. These dried coverslips were coated with 2 mg/mL methoxy-poly (ethylene glycol) (mPEG)-silane MW 2,000 and 2 µg/mL biotin-PEG-silane MW 3,400 (Laysan Bio, USA) in 80% ethanol (pH 2.0) and incubated overnight at 70°C. A 40 µm high PDMS mold with 3 inlets and 1 outlet was mechanically clamped onto a PEG-Silane coated coverslip. The chamber was then connected to a Maesflo microfluidic flow-control system (Fluigent, France), rinsed with modified TIRF buffer (regular TIRF buffer supplemented with 50 mM inorganic phosphate: 10 mM imidazole pH 7.4, 34.8 mM K_2_HPO_4_ and 15.2 mM KH_2_PO_4_, 1 mM MgCl_2_, 1 mM EGTA, 0.2 mM ATP, 10 mM DTT, 1 mM DABCO) and incubated with 1% BSA and 10 µg/mL streptavidin in 20 mM HEPES pH 7.5, and 50 mM KCl for 5 min. Capping protein was then anchored on the surface by flowing in 1 nM biotin-SNAP-CP for 5 min. Pre-formed actin filaments (10% Alexa-488 labelled) were then introduced and captured by coverslip-anchored CP at their barbed ends with their distal pointed ends free in solution. These filaments were then exposed to cofilin (and or N-CAP1) in modified TIRF buffer. All experiments were conducted at room temperature.

### Image acquisition and analysis

Single-wavelength time-lapse TIRF imaging was performed on a Nikon-Ti2000 inverted microscope equipped with a 40 mW 488 nm Argon laser, a 60X TIRF-objective with a numerical aperture of 1.49 (Nikon Instruments Inc., USA) and an IXON LIFE 888 EMCCD camera (Andor Ixon, UK). One pixel was equivalent to 144 × 144 nm. Focus was maintained by the Perfect Focus system (Nikon Instruments Inc., Japan). Time-lapsed images were acquired every 10 s using Nikon Elements imaging software (Nikon Instruments Inc., Japan).

Images were analyzed in Fiji (Schindelin et al., 2012). Background subtraction was conducted using the rolling ball background subtraction algorithm (ball radius 5 pixels). For each condition, between 50 and 100 filaments were acquired across multiple fields of view. To determine the rate of depolymerization, the in-built kymograph plugin was used to draw kymographs of individual filaments. The kymograph slope was used to calculate pointed-end depolymerization rate of each individual filament (assuming one actin subunit contributes 2.7 nm to filament length). Slopes were only measured for long regions of kymographs not showing a visible pause in depolymerization (presumably due to photodimerization or due to surface interactions). Data analysis and curve fitting were carried out in Microcal Origin. Depolymerization rate (*D_p_*) in figures 4b and 4d were fit to the following expression (Ankita et al., 2023):

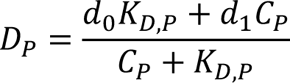

where *d_0_* is the depolymerization rate in absence of N-CAP (determined experimentally), and *d_1_* is the depolymerization rate at N-CAP concentration *C_P_* and *K*_D_ is the dissociation constant of N-CAP. All experiments were performed at least three times and yielded similar results. The data shown are from one experiment each.

### Co-sedimentation Assays

Actin filaments were polymerized by addition of 50 mM KCl, 1 mM MgCl_2_ and 0.2 mM EGTA. 5 µM F-actin was incubated with a range of concentrations of cofilin either in a buffer without 50 mM P_i_ (10 mM imidazole pH 7.4, 10 mM DTT, 0.2 mM ATP, 50 mM KCl, 1 mM MgCl_2_ and 1 mM EGTA) or with 50 mM P_i_ (10 mM imidazole pH 7.4, 34.8 mM K_2_HPO_4_ and 15.2 mM KH_2_PO_4_, 1 mM MgCl_2_, 1 mM EGTA, 0.2 mM ATP and 10 mM DTT) for 15 minutes at room temperature. Then samples were ultracentrifuged for 20 mins at 450,000 × *g.* Supernatant from each sample was discarded and the pellets were washed twice with F-buffer and then resuspended in 100 µl of G-buffer. Samples were run on a gel, and a standard curve with known cofilin concentrations was used to determine the concentration of cofilin-1 in the pellets.

## Acknowledgement

We thank Prof. Shoichiro Ono for advice on co-sedimentation assays.

## Author contributions

E.T., B.M. and S.S. conducted experiments and analyzed data. H.U., B.M.

H.U. and S.S. prepared key reagents. E.T. and S.S. prepared figures. S.S. designed experiments and supervised the project. S.S. and E.T. wrote the first draft of the manuscript and all authors contributed to the editing. S.S. acquired funding.

